# NON-LINEAR SUSCEPTIBILITY TO INTERFERENCES IN DECLARATIVE MEMORY FORMATION

**DOI:** 10.1101/2021.06.29.450433

**Authors:** D. Moyano Malen, Carbonari Giulia, Bonilla Matías, E. Pedreira María, I. Brusco Luis, Kaczer Laura, Forcato Cecilia

## Abstract

After encoding, memories go through a labile state followed by a stabilization process known as consolidation. Once consolidated they can enter a new labile state after the presentation of a reminder of the original memory, followed by a period of restabilization (reconsolidation). During these periods of lability the memory traces can be modified. Currently, there are studies that show a rapid stabilization after 30 min, while others show that stabilization occurs after longer periods (e.g. 6 h). Here we investigate the effect of an interference treatment on declarative memory consolidation, comparing distinct time intervals after acquisition. On day 1, participants learned a list of non-syllable pairs (List 1). Immediately after, 30 min, 3 h or 8 h later, they received an interference list (List 2) that acted as an amnesic agent. On day 2 (48 h after training) participants had to recall List 1 first, followed by List 2. We found that the List 1 memory was susceptible to interference when the List 2 was administered immediately or 3 h after learning; however, shortly after acquisition (e.g. 30 min) the List 1 memory becomes transiently protected against interference. We propose the possibility that this rapid memory protection could be induced by a fast and transient neocortical integration (where the memory is transiently protected) becoming partially independent from the hippocampus followed by a hippocampal re-engagement where the memory becomes susceptible to interferences again. Our results open a discussion about the contribution of molecular and systemic aspects to memory consolidation.

## 1. INTRODUCTION

Memory consolidation encompasses different processes at multiple levels of organization and function in the brain, from the molecular to the behavioral level, and over a temporal spectrum ranging from seconds to months and years that transform, stabilize and update memory traces according to contextual demands (Dudai et al., 2015). Consolidation involves modifications of the synapses concerning the engram (synaptic consolidation) as well as a redistribution of information to long-term storage areas (system consolidation) (Dudai et al., 2015). However, once consolidated, memories are not fixed. After the presentation of a cue associated with the original information (reminder), stored memories can return to a labile state followed by a process of re-stabilization (reconsolidation), dependent on protein synthesis and gene expression (Nader et al., 2000). Even though reconsolidation is not a recapitulation of the consolidation process, they share common molecular mechanisms (Alberini, 2005). Furthermore, during both labile periods (i.e. after acquisition or by reminder presentation) memories can be modified: impaired, strengthened or updated in content (Dudai, 2004, 2012; Haubrich & Nader, 2016). Regarding the time windows of lability, several studies support the idea that once stabilization/re-stabilization is accomplished, memories can no longer be modified without external reactivation (Nader et al., 2000, Forcato et al., 2007). However, other studies propose that memory formation involves multiple waves of consolidation processes even without external reactivation of the memory trace (Alberini, 2005). Therefore, in the present study we aimed to bridge these findings using the lability to interference as a tool to address the dynamics of memory stabilization.

The interval in which memory is labile and can be impaired have been studied in different animal models and also in humans, on different types of memory using different types of amnesic agents (Duncan,1949; Gerard, 1995; Thompson & Dean, 1995; Heriot & Coleman, 1962; Agranoff et al.,1966; Nader et al., 2000; Forcato et al., 2007; Muellbacher et al., 2002). The first experiments about memory consolidation in animal models were carried out using electroconvulsive shock (ECS) as interference: the ECS was administered in different time points after learning (Duncan,1949; Gerard, 1995; Thompson & Dean, 1995; Heriot & Coleman, 1962). Duncan (1949) using different intervals between the shuttle box training and the ECS on rats (20 sec, 40 sec, 4 min, 15 min, 1 h, 4 hand 14 h) found that if 1 h or more elapsed between the end of a training and the ECS, there was no apparent memory loss, suggesting that 1 h after learning, the memory can no longer be interfered. By the same time, Agranoff et al. (1966) working on shuttle-box learning in goldfish, showed the time window in which memory was sensitive to interference using puromycin, a protein synthesis inhibitor. The protein synthesis inhibitor was administered to separate groups at 0, 30, 60 and 90 min after training. The sensitivity of memory to protein synthesis inhibition was over by about one hour. In humans, Müller and Pilzecker (1900) in their pioneering study found that the memory of a recently acquired list of syllables can be disrupted if the subject has to learn a second list of syllable pairs in short succession. In contrast, the two lists of syllables were well remembered if they were presented spaced by 2-3 h. Other studies using a motor learning and a second task as interference in humans showed that 6 h after learning, the memory was no longer susceptible against interferences (Shadmehr & Holcomb, 1997; Walker et al., 2003). Nevertheless, in a recent study, Kaczer and colleagues (Kaczer et al., 2018) using a new-word learning paradigm, observed that the interference (i.e. other set of new words) affected the memory consolidation only when it occurred 5 min after training, but this effect was not observed when the interference was presented at 30 min, 4 or 24 h after. A question that emerges from these studies is whether the resistance to interference is a gradually emergent property after acquisition, or instead if there are multiple time windows that could be revealed.

Furthermore, consolidation and reconsolidation not only share similar molecular mechanisms (Alberini, 2005) but also similar time windows in which memory is sensitive to interferences (after learning or after the presentation of the reminder). Studies in animal models using ECS or protein synthesis inhibitor as an amnestic agent, showed that memory can be interfered with up to 6 h after labilization (Schneider & Sherman, 1968; Nader et al., 2000). In humans, Forcato et al. (2007) showed that a consolidated declarative memory (list of syllable pairs, List 1) could be reactivated by the presentation of a reminder and a second learning session (of another list of syllable pairs, List 2) could interfere with its re-stabilization when it was presented 5 min or 6 h after the reminder but not 10h later when the memory was already re-stabilized. However, Shen et al., (2019) showed a rapid re-stabilization after 30 min, in the same line of the findings by Kaczer et al. (2018). They observed that presenting the interference immediately or 20 minutes after reactivation interrupted memory re-stabilization, while if it was presented 30 or 40 minutes after, it had no effect (Shen et al., 2019). However, they did not evaluate the interference effect at longer intervals (e.g. 1 h and 6 h), leaving the possibility that a second window of lability could be observed.

In summary, there seems to be a consensus about the susceptibility to interference on memory consolidation/reconsolidation when the amnesic agent is administered immediately after training/reactivation (Müller & Pilzecker, 1900, Walker et al., 2003, Forcato et al., 2007, Shen et al., 2019, Kaczer et al., 2018). Furthermore, this time window of lability has been proposed to be closed at long delays, such as > 6 h or 24 h, when no interference effect is observed neither in consolidation nor reconsolidation processes (Nader et al., 2000, Forcato et al., 2007, Walker et al., 2003, Herszage & Censor, 2017). However, there are findings that point to the existence of a second time window of memory protection against interferences, around 30-40 min after acquisition (Kaczer et al., 2018, Shen et al., 2019).

Taking all these in mind, we aimed to study the effect of interference treatment after short and long periods after acquisition, on declarative memory consolidation. With this aim, we performed a two-day experiment with six groups. Participants learned a list of five pairs of non-syllable pairs on day 1 (List 1). Immediately after, 30 min, 3 h or 8 h later they received an interference list (List 2) that acted as an amnesic agent. They were finally tested on both lists on day 2 (48 h after training).

## 2. MATERIALS AND METHODS

### 2.1. Participants

192 volunteers were enrolled in the study (145 women, 47 men). The participants were undergraduate and graduate students with ages ranging from 18 to 36. They were recruited via mail and social media pages from our laboratory (Twitter, Facebook and Instagram). In order to be able to participate, they first had an interview with the experimenter who explained the procedures and next they had to tick a box in an online consent form approved by the *“Comité de Ética en Investigación Biomédica del Instituto Alberto C. Taquini”*, in accordance to the principles expressed in the Declaration of Helsinki. Among the participants that concluded the experiment, three gift cards from a bookstore were raffled and the winner was informed in the social media pages and by mail.

None of the participants reported ongoing medication, health problems, medical interventions, or history of psychiatric, neurological, or sleep disorders.

Subjects that reached at least 55% of correct responses during the last four training trials (11/20 correct responses) were included in the analysis. 71 subjects were excluded from the analysis because they did not reach the learning criterion (41), did not followed/understand the instructions of the task (15), wrote the syllables in a piece of paper during the training session or the retention interval (9), used psychopharmaceuticals that they had not reported it the initial interview (3), slept a nap during retention interval (1) or did not respected the indicated schedule for carry out the experiment (2). The final sample included 121 participants (89 women), with ages ranging from 18 to 36 (25.74 ± 0.45 years).

### 2.2. Experimental groups

Participants were randomly assigned to one of six groups. All groups performed the List 1 training on day 1 and were tested on day 2 (Fig. 1A). The groups differed in the moment they received the interference task on day 1. The “G-5min” group (n = 23) received the interference task immediately after the L1-training. The “G-30 min” group (n = 20) received the interference task 30 min after the L1-training. The “G-3h” group (n = 18) learned the List 1, and received the interference task after 3 h. The “G-8h” group (n = 17) received the interference task 8 h after the L1-training. Two more control groups were assessed. The control List 1 group (“CTL-L1”, n = 22) received the List 1 training on day 1 but did not learn the interference task. Participants in the control List 2 group (“CTL-L2”, n = 21) learned only the interference task on day 1 and were tested on day 3.

**Fig. 1.**
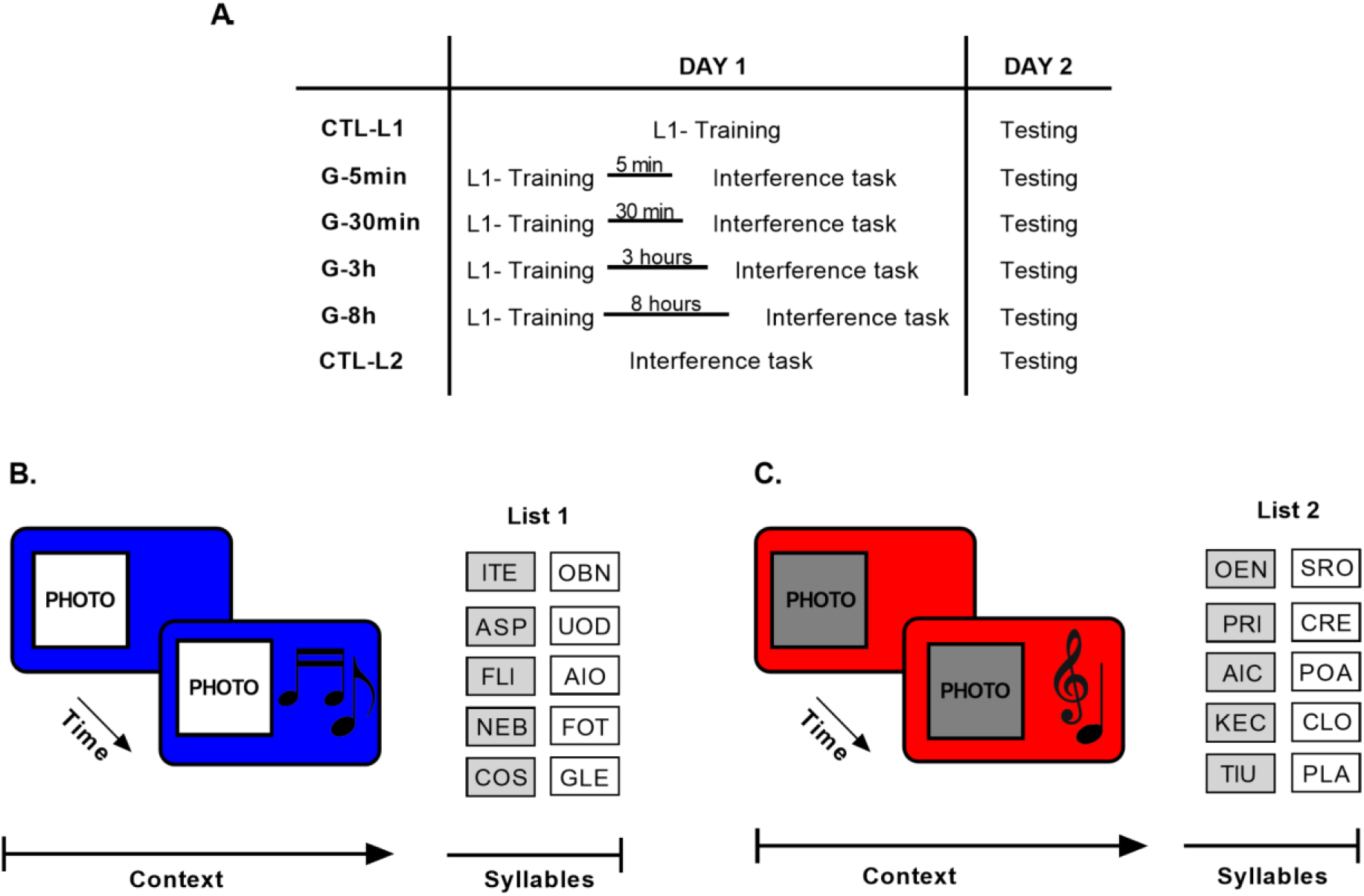
Experimental design and memory task. **A. Protocol**. The experiment was run on two days, 48 h apart. On day 1, all the experimental groups received the List 1 training and were tested on day 2 (first for List 1 and after that for List 2). The groups differed in the moment they received the interference task (5 min, 30 min, 3 h or 8 h after List 1 training). The “CTL-L1” and “CTL-L2” groups were only trained and tested in one list (List 1 and List 2, respectively). **B. L1 Training**. The training session consisted of 10 trials. Each trial started with the presentation of the blue background and image of an Italian coast for 4 sec, followed by the same stimuli accompanied by the tarantella music for another 4 sec. After that, the five pairs of cue-response syllables (List 1) were presented successively and in random order. **C. Interference task**. The training of the interfering task was the same as in B but with a different context (red background colour, the image of a forest and classical music) and different five pairs cue-response syllables (List 2).

### 2.3. Experimental design

The experiment was conducted online due to the Covid-19 pandemic. We used Gorilla Experiment Builder (www.gorilla.sc) to create and host our experiment (Anwyl-Irvine, Massonnié, Flitton, Kirkham & Evershed, 2018). It was a semi-directed experiment where people interested in participating had a first interview with the experimenter. Once the participants agreed to participate, they received the link to access the platform.

The participants could start the experiment between 9:00 h and 18:00 h. Only “G-3h” and “G-8h” groups had more specific indications regarding the start time of the experiment: the “G-3h” group started the experiment between 9:00 h and 16:00 h and the “G-8hs” group started the experiment between 8:00 h and 10:30 h. Thus, the interference task was not presented at night to avoid placing it near the onset of sleep. In all experimental groups, the participants gave their consent, filled out a personal data questionnaire and completed the psychological tests and questionnaires (Beck Depression Inventory II, State-Trait Anxiety Inventory, Pittsburgh Sleep Quality Index and Morningness-Eveningness Questionnaire). After that, they received the instructions for the memory task and completed a demo test. Then the procedure changed for each group.

The “G-5min” group learned the first list of syllable-pairs (List 1) and immediately after learned the interference task (List 2) between 9:00 h and 18: 00 h (14:51 ± 0.02 h). The “G-30min” group learned the first list of syllable-pairs (List 1) at any time between 9:00 h and 18:00 h (13:56 ± 0.09 h). After 30 minutes of completing this first part, the participants received an email to re-enter the platform and complete the second part of Day 1: the interference task. The “G-3h” group followed the same procedure as the “G-30min” group, except that they had a 3 h period between the List 1 and the interference task, thus they started the experiment at any time between 9:00h and 16:00h (13: 52 ± 0.09 h). The “G-8h” group learned the first list of syllable-pairs (List 1) between 8:00h and 10:30 h (9:29 ± 0.03 h). They had an 8 h period between the List 1 and the interference task. Both the “G-3h” group and the “G-8h” group continued with their normal activities during the 3 h or 8 h, respectively. Moreover, they were instructed not to sleep during this period. The procedures for the control groups were similar to the experimental groups but they only learned one list of syllables. The “CTL-L1” group learned the List 1 and the “CTL-L2” group learned the List 2 at any time between 9:00 h and 18:00 h (13:41 ± 0.10 h and 14:20 ± 0.13 h, respectively).

On day 2, at 48 h on day 1, all groups received an email at 9:00 h with the link to re-enter the platform and performed the last session of the experiment. They could enter the link at any time between 9:00 h and 18:00 h (G-5min: 15:19 ± 0.02 h; G-30 min: 13:45 ± 0.10 h; G-3h: 13:49 ± 0.11 h; G-8h: 13:07 ± 0.09 h, CTL-L1: 14: 16 ± 0.12 h, CTL-L2: 15: 26 ± 0.11 h). The experimental groups were tested for retrieval on both lists: first for List 1 and then for List 2, while the control groups were only tested on the single list.

### 2.4. The task

The task consisted of memorizing five pairs of nonsense syllables, associated with a context formed by a background color on the computer screen, an image and music, presented through headphones (Moyano et al., 2019). The syllables were formed by three letters (Fig. 1B and Fig. 1C). We have previously shown that including a context associated to the list of syllables improved memory retention (Forcato et al., 2007).

#### 2.4.1. List 1 training session

List 1 was constituted by five pairs of nonsense cue-response syllables: **ITE**-OBN, **ASP**-UOD, **FLI**-AIO, **NEB**-FOT, **COS**-GLE (bold type: cue-syllable; regular type: response-syllable, Fig. 1B). The training session of List 1 consisted of 10 trials, associated to a context consisting of a blue background color, an image of an Italian coast and a tarantella melody (Fig. 1B). The first trial started with the presentation of the context: first, the image of the Italian coast with blue background color appeared on the screen for 4 sec. Then, a tarantella melody played along with the image and background color for another 4 sec. After that, the context continued while the syllables were presented. First, one cue-syllable appeared at the left top side of the monitor’s screen and an empty response box was displayed on the right top. Next, the corresponding response-syllable appeared in the response box and stayed there for 4 sec. Immediately thereafter, the syllable pair disappeared and another cue-syllable was shown one line below and the process was repeated until the list was complete. Each cue-syllable was taken at random and successively from the list of five until the trial was complete. Thus, in the first trial the participants observed how all five pairs were completed once and in the successive nine trials subjects were required to write down the corresponding response-syllable for each cue-syllable presented. Each of those nine trials began with the presentation of one cue-syllable at the left top side of the monitor’s screen and an empty response box on the right top. Subjects were given 5 sec to write the corresponding response-syllable. Once that period had finished, three situations were possible: first, if no syllable was written down, the correct one was shown for 4 sec; second, if an incorrect syllable was written, it was replaced by the correct one and it was shown for 4 sec; and third, if the correct response was given, it stayed for a further 4 sec. Immediately thereafter, the syllable pair disappeared and another cue-syllable was shown one line below and the process was repeated until the list was complete. After the whole list was presented, a black background was shown for 3 sec and the procedure was repeated until the entire list of syllable-pairs was completed nine times. The List 1 training session took about 10 min.

#### 2.4.2. Interference task

The interference task consisted of the learning of another list of syllable pairs, List 2, which was formed by five different pairs of nonsense cue-response syllables: **OEN**-SRO, **DRI**-CRE, **AIC**-POA, **TIU**-PLA, **KEC**-CLO (Fig. 1C). Learning of the interference task was similar to the List 1 training session, but with a different context (red background colour, the image of a forest and classical music). Like the List 1 training session, it was formed by 10 trials. In the first trial, participants observed how the five syllable pairs were completed once and in the successive nine trials they had to write down the corresponding response-syllables. Feedback procedures were the same as for List 1 training. The interference task took about 10 min.

#### 2.4.3. Testing session

On day 2, List 1 and List 2 memory was tested. List 1 memory was always tested first before List 2 memory. For both lists, testing was formed by four trials each, similar to the training session, i.e. including recall of each of the syllable pairs twice. Cue-syllables were taken at random and successively from the list of five. Subjects were required to write down the corresponding response-syllable within 5 sec. The testing session took about 4 min (2 min per List).

The written responses were registered and an error was considered when a participant wrote an incorrect response-syllable or if they did not answer at all. Furthermore, we classified the errors into four categories: “void”, when no response was written down; “intralist”, when a wrong response-syllable from the same list was written down; “intrusion”, when a syllable from the other list was written down; and “confusion”, when the written response-syllable was not included in any of the lists.

#### 2.4.4. Demo

Before the List 1 training session, participants were presented with a demo program to receive all the instructions and to make sure that all participants understood the task. The demo program consisted of two trials, similar in structure as the training session but with another context and two different pairs of nonsense syllables.

### 2.5. Method for evaluating the impairment on the L1 consolidation process

A common way to reveal the presence of the consolidation process, is to present an amnesic agent after learning to interfere with the stabilization of the memory trace (Forcato et al., 2007). Thus, the presence of such process is revealed by the absence (impairment) of the memory at testing session. However, this direct method of evaluation may be sometimes misleading or inapplicable given the fact that memories are not stored in isolation from other memories but integrated into complex associative networks (Levy & Anderson 2002; Berman et al. 2003; Debiec et al. 2006), and then the activation of one memory may interfere with the retrieval of interest (McGeoch 1932; Postman 1971; Anderson & Neely 1996). In other words, a faulty retrieval at testing may be due to either problems in encoding storage or simultaneous retrieval of related information (Mayes & Downes 1997). Forcato et al. (2007) proposed that when working with two related memories, the direct method would lack specificity (Forcato et al., 2007). Given the existing interaction between consolidated memories, they proposed the use of an indirect method based on a temporal “forgetting” effect. This effect, termed retrieval-induced forgetting (RIF) (Anderson et al.1994; MacLeod & Macrae 2001), shows that retrieval of target memories could temporarily block subsequent retrieval of other, related memories. This RIF effect is only observed when the memory trace that is recalled first is intact, as a consequence, a poor performance was expected for the second task at testing. Whereas the RIF effect is not observed when the first recalled memory trace is impaired, and therefore the absence of RIF might become a good indicator of a defective target memory (evidenced by no RIF effect; Forcato et al., 2007, 2009, 2013).

### 2.6. Tests and questionnaires

#### Beck Depression Inventory II (BDI-II)

The BDI-II (Beck, Steer & Brown, 1996) is a 21 question self-report inventory, assessing the somatic, cognitive and affective symptoms of depression in the preceding 2 weeks. Each question has a set of at least four possible responses, ranging in intensity. When the test is scored, a value of 0 to 3 is assigned for each answer and then the total score is compared to a key to determine the depression’s severity. Higher scores indicate depressive symptoms being more severe.

#### State-Trait Anxiety Inventory (STAI)

The STAI (Spielberger et al., 1983) is a self-reported questionnaire composed by 40 items developed with the aim of evaluating two different types of anxiety: state anxiety (emotional condition transitory), whose reference frame is the “now, at this moment” (20 items), and the anxiety trait (anxiety tendency relatively stable), whose reference frame is “in general, in most of the times”. The STAI has a Likert-type response format with four options (0=almost never/nothing; 1=some/some times; 2=quite/ often; 3=a lot/almost always).

#### Pittsburgh sleep quality index (PSQI)

The PSQI (Buysse et al., 1989) measures a broad range of symptoms of sleep disturbances over a 1-month period. This is composed of 19 questions which reflect seven major components. All seven components are then summed up to create a scale from 0–21 points. Higher scores indicate a worse sleep quality.

#### Morningness-Eveningness Questionnaire (MEQ)

The MEQ is a widely-used international questionnaire validated by Horne and Ostberg (Horne & Ostberg, 1976). This questionnaire is composed of 19 items. It is self-administered and measures the person’s peak alertness/sleepiness (morning or evening). MEQ consists of 19 questions allowing to calculate a total score between 16 and 86; scores ≤30 indicate definite evening type, 31–41 indicate moderate evening type, 42–58 intermediate type, 59–69 moderate morning type, and 70–86 definite morning type.

### 2.7. Data analysis and statistics

Statistical analysis was done with SPSS version 25 (IBM Corporation). We calculated the level of learning as the mean total number of correct responses in the four last training-trials. The level of learning was analyzed with one Way-ANOVA with “group” as a between subjects factor followed by Bonferroni Post-hoc comparisons. We calculated the memory change for each List as the number of correct responses in the first trial at testing on day 2 minus the number of correct responses in the last trial of training on day 1. Thus, positive values mean memory gain and negative values, memory loss. To determine if there was a significant decrease in memory from day 1 to 2, we further performed two tailed one sample t-test for each group compared to the value zero (no memory change). The memory change for each List was analyzed with one-way ANOVA with “group” as a between subjects factor followed by Bonferroni Post-hoc comparisons. Comparisons were made against the control group of each list, since it is the group that was only trained and tested on one list.

For the analysis of type of errors we calculated the change of error’
ss type for: “void”, “intralist”, “confusion” and “intrusion” errors from each List. The change of error’s type was defined as the percentage of errors in the last trial at training on day 1 minus the percentage of the errors in the first trial of testing on day 3. Thus, negative values mean memory loss, i.e. more percentage of a certain type of error at testing. The change of error’s type was analyzed with independent one-way ANOVAs. For the “intrusion” errors we excluded the control group for each list, because it only learns one of the two lists, therefore it does not present this type of error. For all analyses, we applied a significant threshold of p = 0.05.

We further analyzed the State Anxiety, Trait Anxiety, BDI-II, PSQI and MEQ in all conditions with one-way ANOVA with “group” as a between subjects factor.

In order to analyze the dynamics of the RIF effect on List 2 memory change, all time points (0 min, 30 min, 3 h, 8 h) were fitted to a cubic function [*f*(*x*) = *a* ∗ *x*3 + *b* ∗ *x* 2 + *c* ∗ *x* + *d*] using a cubic spline interpolation in Python. Also, to evaluate the dynamics of the interference task on the List 1 memory change, all time points were fitted to another cubic spline function.

## 3. Results

### 3.1. L2-List performance

As in previous studies we observed that a faulty retrieval at List 1 testing could be due to storage impairment or to simultaneous retrieval interferences (Forcato et al., 2007), we here used the RIF effect on L2 testing to reveal the impairment in the consolidation of L1. That is, the act of remembering could temporarily block the subsequent retrieval of related information (RIF effect, Anderson et al., 1994). Thus, if the L1 memory is intact, its retrieval may interfere with subsequent retrieval of L2 memory. Otherwise, when L1 memory is impaired, its retrieval will not interfere with the retrieval of related memories (No-RIF) (Forcato et al., 2007; 2013).

The groups reached significantly different levels of List 2 learning in the last four training trials (Fig. 2.A1, “CTL-L2”: 79.52 ± 2.88 %, “G-5min”: 88.91 ± 2.20 %, “G-30min”: 92.75 ± 2.68 %, “G-3h”: 86.39 ± 3.59 %, “G-8h”: 83.24 ± 3.03 %, one way ANOVA F(4,98) = 3.38, p = 0.013). Specifically, the “G-30 min” group showed a better performance than the “CTL-L2” group (Bonferroni p = 0.01). No significant differences were found between the “CTL-L2” and the other experimental groups (all p > 0.16). Thus, in order to study the time window of lability of the declarative memory consolidation, we calculated the memory change (the number of correct responses at the first testing trial minus the number of correct responses at the last training trial).

**Fig. 2.**
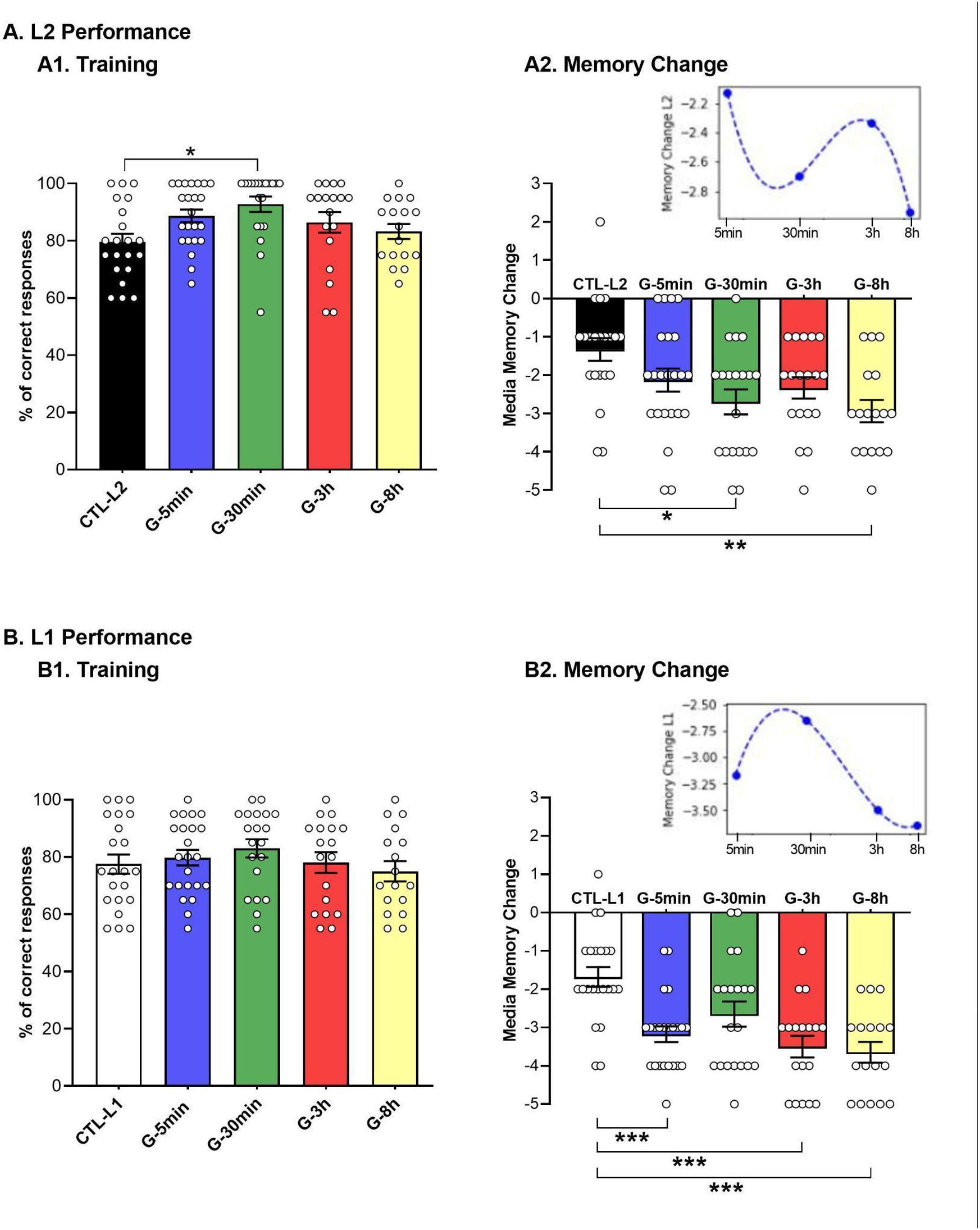
Declarative consolidation dynamics. **A. List 2 performance. A1. Training.** The mean total number of List 2 correct responses in the four last training trials ± SEM. **A2. List 2 memory change**. Mean memory change (number of correct responses at the first List 2 testing trial minus the number of correct responses at the last List 2 training trial) ± SEM. *Inset:* List 2 mean memory change was fitted to a cubic function: Time is displayed on a logarithmic scale. **B. L1 List Performance. A1. Training**. The mean total number of List 1 correct responses in the four last training trials ± SEM. **A2. List 1 memory change**. Memory change mean ± SEM. *Inset:* List 1 mean memory change was fitted to a cubic function: Time is displayed on a logarithmic scale. * p < 0.05; **, p < 0.01; ***, p < 0.001.

We first observed that the performance of all groups significantly decayed between training and testing (T-test: “CTL-L2” T_20_ = -4.51 p < 0.001, “G-5min” T_22_ = - 7.02 p < 0.001, “G-30min” T_19_ = -8.30 p < 0.001, “G-3h” T_17_ = -8.33 p < 0.001, “G-8h” T_16_ = -10.13 p < 0.001). Furthermore, the groups significantly differed in the L2 memory change (Fig. 2A2, CTL-L2: -1.33 ± 0.30, “G-5min”: -2.13 ± 0.30, “G-30min”: -2.70 ± 0.33, “G-3h”: -2.33 ± 0.28, “G-8h”: -2.94 ± 0.29, one-way ANOVA, F(4,98) = 4.17, p = 0.004). The “G-5min” and “G-3h” groups showed a similar decay than the “CTL-L2” group that was only trained and tested on List 2 (Bonferroni p = 0.53 and p = 0.23, respectively), evidencing no RIF effect, whereas the “G-8h” group showed a significant higher decay in memory change than the “CTL-L2” group, evidencing an intact RIF effect (Bonferroni p = 0.004). Participants in the “G-30min” group had significantly more decay than the “CTL-L2” group (Bonferroni p = 0.016), showing an intact RIF effect (Fig. 2A2). It is important to highlight that the “G-30min” and the “G-8h” groups behaved the same related to “CTL-L2” group. There were no significant differences on L2 memory change between the experimental groups (all ps > 0.63). Thus, the interference task (List 2) presented immediately or 3 h after List 1 training interfered with the stabilization of the List 1 memory (evidenced by no RIF effect on List 2). However, when the interference task was presented 8 h later, or 30 min after training it did not impair List 1 memory stabilization (evidenced by an intact RIF effect).

An analysis of the dynamics of List 2 memory change revealed that the curve (*f*(*x*) = −0.97 ∗ *x* 3 + 1.14 ∗ *x* 2 − 4.10 ∗ *x* + 1.94) presented two minimums (higher List 2 memory decay), one at 30 min and the other at 8 h, evidencing higher RIF effect.

### 3.2. L1 List performance

There were no significant differences between groups in the level of List 1 learning (Fig. 2B1, “CTL-L1”: 77.50 ± 3.33 %, “G-5min”: 79.78 ± 2.71 %, “G-30min”: 83.00 ± 3.17 %, “G-3h”: 78.06 ± 3.62 %, “G-8h”: 75.00 ± 3.79 %; F(4,99) = 0.79, p = 0.53). Furthermore, when examining the memory change of List 1 all groups showed a significant decay in performance between training and testing (T-test: “CTL-L1” T21= -6.51 p < 0.001, “G-5min” T_22_ = -15.47 p < 0.001, “G-30min” T19 = -8.11 p < 0.001, “G-3h” T17 = -12.37 p < 0.001, “G-8h” T16 = -13.49 p < 0.001). There were significant differences between groups at memory change (Fig. 2B2, “CTL-L1”: -1.70 ± 0.26, “G-5min”: -3.17 ± 0.21, “G-30min”: -2.65 ± 0.33, “G-3h”: -3.5 ± 0.63, “G-8h”: -3.65 ± 0.27, one way ANOVA, F(4,99) = 8.93 p < 0.001). Specifically, the “G-5min”, “G-3h” and “G-8h” groups showed a significant higher decay than the “CTL-L1” group that was only trained and tested on List 1(Bonferroni, p = 0.001, p < 0.001 and p < 0.001, respectively). The “G-30min” and the “CTL-L1” groups did not significantly differ (Bonferroni p = 0.10). There were no significant differences on List 1 memory change between the experimental groups (all ps > 0.13).

Furthermore, an analysis of the dynamics of List 1 memory change showed that the curve (*f*(*x*) = 0.07 ∗ *x* 3 − 0.95 ∗ *x* 2 + 3.65 ∗ *x* − 6.87) presented a maximum at 30 min, evidencing a time point where the memory is more protected against simultaneous interferences.

### 3.3. Type of errors

We analyzed the change in error’
ss types for each list (Table 1). For List 2 there was a significant difference between groups for the “void” type errors (“CTL-L2”: -12.38 ± 4.25 %, “G-5min”: -28.70 ± 5.31%, “G-30min”: -39.00 ± 6.24 %, “G-3h”: -24.44 ± 5.95 %, “G-8h”: -31.76 ± 7.49 %, one way ANOVA, F (4,98) = 2.97, p = 0.023). We observed that the “G-30min” showed a significantly higher decay than the “CTL-L2’’ (Bonferroni p = 0.014). However, there were no significant differences for the “G-5min”, “G-3h” and the “G-8h” compared to the “CTL-L2” group (p = 0.39 p = 1 and p = 0.24, respectively). There were no significant differences between groups neither for the confusion, nor for the intralist and intrusion type of errors (One way ANOVAs, F(4,98) = 1.80, p = 0.14, F(4,98) =1.30, p= 0.28, F(3,77) = 1.99, p = 0.12, respectively). For List 1, there was a significant difference between groups for the “void” type errors (“CTL-L1”: -15.45 ± 4.15 %, “G-5min”: -33.91 ± 5.54 %, “G-30min”: -40.00 ± 6.32%, “G-3h”: -28.89 ± 6.71 %, “G-8h”: -42.35 ± 8.02 %, one way ANOVA, F(4,99) = 3.18, p = 0.02). Specifically, the “G-30min” and “G-8h” groups showed higher decay of “void” type errors that “CTL-L1” (Bonferroni p = 0.043 and p = 0.028, respectively). However, no significant differences were found for the “G-5min” and “G-3h” groups compared to the “CTL-L1” (Bonferroni all ps > 0.25). There were no significant differences between groups neither for the confusion nor for the intralist and intrusion type of errors (One way ANOVAs, F(4,99) = 1.87, p = 0.12, F(4,99) = 0.83, p = 0.51, F(3,77) = 2.05, p = 0.11, respectively).

**TABLE 1.** Type of errors.

|  | Void | Confusion | Intralist | Intrusion |
| --- | --- | --- | --- | --- |
| **List 2** |  |  |  |  |
| **CTL-L2** | -12.38 ± 4.25 | -12.38 ± 4.25 | -2.86 ± 4.64 | - |
| **G-5min** | -28.70 ± 5.31 | -4.35 ± 3.55 | -6.96 ± 2.98 | -2.61 ± 1.44 |
| **G-30min** | -39.00 ± 6.24 | -15.00 ± 4.56 | 0.00 ± 2.05 | 0.00 ± 0.00 |
| **G-3h** | -24.44 ± 5.95 | -13.33 ± 3.62 | -6.67 ± 2.80 | -2.22 ± 2.22 |
| **G-8h** | -31.76 ± 7.49 | -21.18 ± 7.17 | 1.18 ± 2.70 | -5.88 ± 2.28 |
| **List 1** |  |  |  |  |
| **CTL-L1** | -15.45 ± 4.15 | -16.36 ± 3.64 | -0.91 ± 2.07 | - |
| **G-5min** | -33.91 ± 5.54 | -19.13 ± 3.44 | 1.74 ± 3.31 | -12.17 ± 3.26 |
| **G-30min** | -40.00 ± 6.32 | -18.00 ± 5.01 | 6.00 ± 4.13 | -1.00 ± 1.00 |
| **G-3h** | -28.89 ± 6.71 | -31.11 ± 4.03 | -2.22 ± 3.19 | -7.78 ± 3.29 |
| **G-8h** | -42.35 ± 8.02 | -25.53 ± 5.21 | 1.18 ± 4.36 | -8.24 ± 5.16 |

Mean percentage memory change of error's type for each List ± SEM.

### 3.4. Emotional variables and Questionnaires

There were no significant differences between groups at STAI State Anxiety (F(5,120) = 0.55, p = 0.74), STAI Trait Anxiety (F(5,120) = 1.35, p = 0.25), BDI-II (F(5,120) = 0.76, p = 0.58), PSQI (F(5,120) = 0.95, p = 0.45), and MEQ (F(5,120) = 0.49, p = 0.78) (Table 2).

**TABLE 2.**
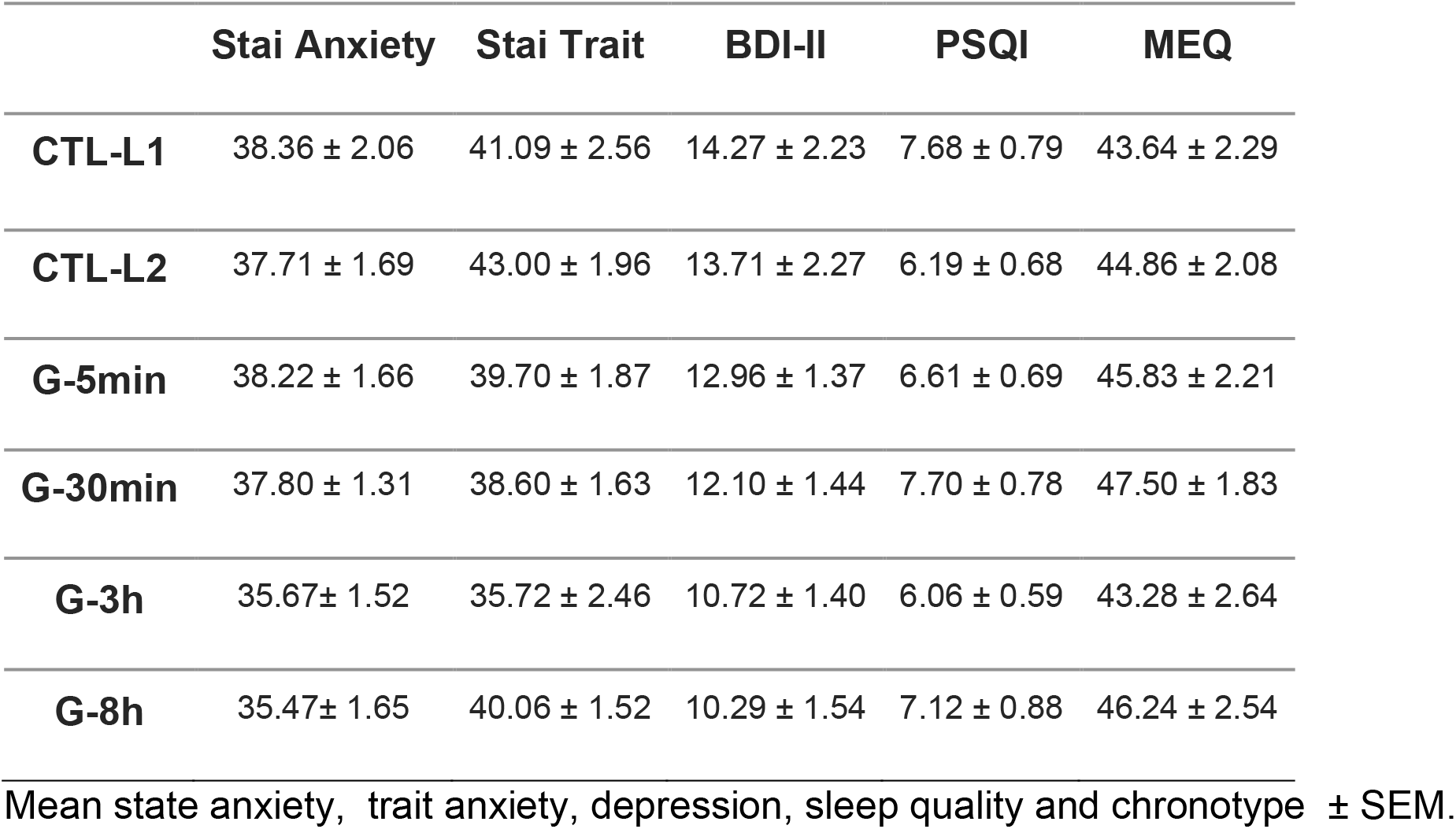
Emotional variables and questionnaires.

|  | Stai Anxiety | Stai Trait | BDI-II | PSQI | MEQ |
| --- | --- | --- | --- | --- | --- |
| **CTL-L1** | 38.36 ± 2.06 | 41.09 ± 2.56 | 14.27 ± 2.23 | 7.68 ± 0.79 | 43.64 ± 2.29 |
| **CTL-L2** | 37.71 ± 1.69 | 43.00 ± 1.96 | 13.71 ± 2.27 | 6.19 ± 0.68 | 44.86 ± 2.08 |
| **G-5min** | 38.22 ± 1.66 | 39.70 ± 1.87 | 12.96 ± 1.37 | 6.61 ± 0.69 | 45.83 ± 2.21 |
| **G-30min** | 37.80 ± 1.31 | 38.60 ± 1.63 | 12.10 ± 1.44 | 7.70 ± 0.78 | 47.50 ± 1.83 |
| **G-3h** | 35.67 ± 1.52 | 35.72 ± 2.46 | 10.72 ± 1.40 | 6.06 ± 0.59 | 43.28 ± 2.64 |
| **G-8h** | 35.47 ± 1.65 | 40.06 ± 1.52 | 10.29 ± 1.54 | 7.12 ± 0.88 | 46.24 ± 2.54 |

Mean state anxiety, trait anxiety, depression, sleep quality and chronotype ± SEM.

## Discussion

In the present study we showed that the dynamics of declarative memory consolidation seems not to be an all or nothing process. As in previous studies we found that immediately after learning, the memory was labile as well as 3 h after, and could be interfered by a second learning task that acts as an amnesic agent (Forcato et al, 2007; 2009; 2013). However, we have not only replicated previous results showing that after longer periods (> 8 h) the declarative memory was already stabilized (Forcato et al., 2007) but we also found a new time window, shortly after acquisition, where the memory became rapidly protected against interference. That is, we demonstrate that there are at least two time windows susceptible to interference after learning without the experimental induction of memory reactivation.

In order to demonstrate List 1 memory impairment, we used the Retrieval-Induced Forgetting effect (RIF) (Forcato et al., 2007). According to this, the retrieval of a target (L1) memory can temporarily block subsequent retrieval of a related memory (L2) (Anderson et al., 1994; Forcato et al., 2007). We used the presence of RIF on the List 2 List as a tool to reveal the memory impairment on List 1, instead of directly measuring the retrieval of List 1 alone, because we have previously observed that a faulty retrieval of L1 could be due to memory impairment or to simultaneous retrieval of related information (Forcato et al., 2007). Thus, if the List 1 memory storage was intact, its retrieval could interfere with subsequent List 2 memory retrieval (RIF effect). On the other hand, if List 1 memory storage was impaired, its retrieval did not interfere with List 2 retrieval (absence of RIF, Forcato et al., 2007). Hence, we evaluated impairment of List 1 memory by analyzing the presence of the RIF effect on List 2. We observed an intact RIF effect after 8 h as well as after 30 min, indicating successful stabilization of List 1 memory in both conditions. On the contrary, the RIF effect was absent after 5 min and 3 h, suggesting that List 1 memory was not yet stabilized at this time and sensitive to disruption by interference learning. Moreover, these results are supported by the interpolation analysis (Fig. 2A2 *inset*), which evidences two maximum peaks representing the “G-5min” and “G-3h” groups (susceptible to interference) and two minimum peaks representing the “G-30min” and “G-8h” groups (more protected against interference).

In addition to the RIF effect, we also evaluated memory performance of List 1. Considering the previous results showing that the List 1 memory performance at testing of the groups that received an interference task had significantly more errors than the control group, revealing simultaneous retrieval interferences between L1 and L2 memory or List 1 consolidation/reconsolidation impairment, (Forcato et al., 2007; 2009; 2013), we expected a reduction on the List 1 retrieval for all groups.

Nevertheless, contrary to our expectations, the group that received the interference task after 30 min of learning showed a similar performance that the “CTL-L1” group, evidencing differential outcomes for this temporal window not only through the RIF effect but also in the List 1 memory retrieval. Moreover, these results are supported by the interpolation analysis (Fig. 2B2 *inset*), which shows one maximum peak at 30 min after acquisition, indicating that the memory is more protected against interference. Furthermore, when we evaluated the memory change regarding the types of List 1 errors we observed the same pattern of results between the “G-30min” and the “G-8h” groups, that is, a significantly higher decay in the void type errors compared to the control group, suggesting that the List 2 could be interfering with the retrieval of List 1 in the “G-30min” group but to a lesser extent given that the interference is not observe when we evaluated the general performance.

It is interesting to note that this short time window after acquisition, where the declarative memory seems to be transiently protected against interferences, matches to the early consolidation processes that take place within about 30 minutes and induce a fast increase in synaptic strength independent of protein synthesis (Frey & Frey, 2008; Frey & Morris, 1997). However, these early changes are transient and decay after about 90 minutes (Frey & Morris, 1998). So, it is possible to speculate that within about 30 minutes after List 1 acquisition, a rapid stabilization, independent of protein synthesis, occurs protecting the memory against List 2 interference. Nevertheless, this does not explain the results underlying absence of simultaneous retrieval interferences on List 1 for the “G-30min” group, that is, the no significant difference between “G-30min” and “CTL-L1” group as we have previously observed for other time windows (Forcato et al., 2007).

Therefore, we suggest that not only synaptic consolidation would be involved, but also that a rapid system consolidation process could be initiated during learning or shortly after acquisition has ended (Wang & Bukuan, 2015; Brodt et al., 2018). One possibility would be that the List 1 memory is rapidly integrated at the neocortical level, becoming partially independent of the hippocampus, so, in this way protecting it against interference. Brodt et al. (2018) using an object–location association task, observed neocortical plasticity (specifically in the parietal cortex) as early as 1 h after learning and found that it was learning specific. They suggested that new traces are encoded rapidly in the neocortex from the learning onset, challenging traditional models of slow systems consolidation (Brodt et al., 2018). In the same line, studies in rodents have revealed that neocortical cells are already tagged during encoding and have detected experience-dependent microstructural changes as early as 1 h after learning (Kitamura et al., 2017; Xu et al., 2009; Cowansage et al., 2014). However, these studies do not explain the second time window susceptible to interferences that we have found, which could suggest a hippocampal re-engagement. In this line, Alberini (2005) has previously described a model that postulates that memory reconsolidation—i.e., the restabilization of the memory trace that follows the labilization induced by its non reinforced retrieval—could be just a manifestation of a lingering consolidation process. According to this model, consolidation may include a number of subsequent reactivation events whose function is to further strengthen and/or prolong memory retention. This hypothesis predicts that there should be recurrent time windows of susceptibility to consolidation blockers over hours, days, or weeks. Taking into account the present results, we propose that the declarative memories are not protected against interference after a unique time window during the first 24 h after learning, on the contrary, we found that there are at least two time windows of susceptibility. Shen et al., (2019) showed that after memory reactivation a second task could interfere with memory re-stabilization up to 20 min, but no interference was observed after 30 and 40 min proposing that the memory was protected against interference 30 min after acquisition. However, they did not study the effect of interference after longer periods. Thus, considering that consolidation and reconsolidation share similar molecular mechanisms it would be of great interest for the clinical field to study this short time window where the memory is protected against interference. In the last years, lots of studies have been proposing the possibility of using reconsolidation as a therapeutic tool for the intervention of maladaptive memories (Diekelmann & Forcato, 2015; Lee, 2009; Bonilla et al., 2020; Fernandez & Allegri, 2019; Alberini & Ledoux, 2013; Lane, 2015; Soeter & Kindt, 2015; Schwabe et al., 2012). That is, to reactivate a maladaptive memory and to update its content. Our results suggest that the time window where the therapeutic intervention is administered should be carefully taken into account because the time windows of susceptibility seem not to be a linear process, at least regarding memory consolidation. Further experiments should be conducted to understand the consolidation and reconsolidation dynamics.

## Conflict of Interest

The authors declare that the research was conducted in the absence of any commercial or financial relationships that could be construed as a potential conflict of interest.

## Author Contributions

MDM, LK and CF made substantial contributions to the conception and design of the work. LK and CF were responsible for funding acquisition. MDM ran the experiments. MDM and GC performed the data analyses. MDM and GC made the graph art. CF administered and supervised the project. LIB and CF were responsible for resources. MDM, CF and LK contributed by drafting the work. MDM, MB, GC, MEP, LIB, LK and CF contributed to revising it critically.

## Funding

This work was supported by AGENCIA (PICT 2016 N° 0229 to CF).

## Financial Disclosure

The funders had no role in study design, data collection and analysis, decision to publish, or preparation of the manuscript. The authors have declared that no competing interests exist.

## Acknowledgment

We thank Marcelo Fabian Blanco for helping with Gorilla Experiment Builder.

## Notes

### Competing Interest Statement

The authors have declared no competing interest.

